# Evidence that standard metabolic rate and risk-taking to breathe air are linked to boldness, activity level and exploratory behaviour in a catfish

**DOI:** 10.1101/2021.02.02.429387

**Authors:** David J. McKenzie, Thiago C. Belão, Shaun S. Killen, Felipe R. Blasco, Morten B.S. Svendsen, F. Tadeu Rantin

## Abstract

We used an air-breathing catfish, *Clarias gariepinus,* to investigate the hypothesis that individual variation in metabolic rate, and the propensity to take risks to obtain a resource (oxygen from air), would be correlated with behavioural tendencies such as boldness, activity level and exploratory behaviour. The standard metabolic rate (SMR) of 58 juvenile catfish was positively correlated with their rates of aerial respiration in daylight when surfacing was inherently risky. SMR was positively correlated with boldness measured in two contexts, namely the time-lag to resume air-breathing in a potentially dangerous environment (T-res, measured in a respirometer), and the timelag to enter the centre of a novel environment (T-centre, measured in an open field test (OFT)). These two measures of boldness were very highly correlated. Thus, these data support the hypothesis that high SMR and an associated tendency to take risks to acquire resources are linked to increased boldness in animals. Individual SMR was positively correlated with the proportion of time the fish spent moving in the OFT, but was negatively correlated with movement speed. The data confirmed previous observations that these catfish may exhibit a bimodal distribution of T-res phenotypes, whereby individuals either resumed air-breathing relatively rapidly (< 85 min, bold n = 26) or more slowly (> 115 min, shy n = 31) after a startle stimulus. Bold T-res phenotypes had significantly higher SMR than shy; breathed more air during the day, and showed greater boldness but less activity and exploration in the OFT. No parallel bold/shy dichotomy was observed, however, in any measure of boldness in the OFT. Therefore, the data support propositions regarding how SMR and risk-taking should relate to boldness, but provide mixed results about how SMR relates to activity and exploration, and whether bold/shy is a dichotomy or spectrum.

## 1. Introduction

Individuals within animal species exhibit wide and temporally persistent variation in metabolic rates (Burton *et al*., 2011; Killen *et al*., 2016). The mechanisms underlying this variation in energy metabolism are not fully understood but the variation itself is believed to be of ecological significance (Burton *et al*., 2011; Careau & Garland, 2012; Metcalfe *et al*., 2016; Auer *et al*., 2018). In particular, it has been shown that individuals with high metabolic rates (MR) may take more risks to acquire resources (Killen *et al*., 2011, 2012; Careau *et al*., 2015). It has been suggested that this may reflect a life history strategy focussing on rapid growth and maturity, with the trade-off being an increased risk of mortality (Stamps, 2007; Biro & Stamps, 2010; Réale *et al*., 2010; Auer *et al*., 2018). It has also been proposed that high MR phenotypes will exhibit particular behavioural tendencies that are consistent with a high-risk lifestyle; in particular greater boldness but also differences in the tendency to be active and exploratory (Biro & Stamps, 2010; Careau & Garland, 2015; Careau *et al*., 2015; Rupia *et al*., 2016; Krams *et al*., 2017). It has even been suggested that individual variation in energy metabolism may play a role in the evolution of these tendencies (Mathot & Dingemanse, 2015; Sih *et al*., 2015).

At present, there is no consensus about how individual energy metabolism might be linked to tendencies other than boldness, such as activity level and exploration, with various models having been proposed (Careau & Garland, 2012, 2015). The ‘increased intake’ model posits a positive relationship between MR and activity, if the high MR of an individual indicates that it has large metabolic capacity for resource acquisition and processing. The ‘compensation’ model posits a negative relationship, if metabolic capacity is finite and a high MR would constrain the ability of an individual to allocate energy to activity (Careau & Garland, 2012). There are also unknowns about the tendencies themselves. For boldness, it is not clear whether animals are distributed along a continuum of bold to shy, whether there is a clear distinction whereby an individual is either one or the other, or whether this can vary depending on context (Sih *et al*., 2004; Thomson *et al*., 2012; Frost *et al*., 2013; Rupia *et al*., 2016).

Fishes that breathe air provide an excellent model to investigate how individual variation in MR and associated risk-taking to obtain a resource, aerial oxygen, are linked to behavioural tendencies (McKenzie *et al*., 2015; Killen *et al*., 2018; Pineda *et al*., 2020). Air is much richer in oxygen than water and various species of fish have evolved adaptations to rise to the surface and breathe it. They store the air in specialised highly vascularised organs, from which oxygen diffuses into the blood. The fishes all have bimodal respiration, air breathing allows them to supplement oxygen that they obtain from water, by ventilating their gills (Randall *et al*., 1981; Graham, 1997). Air breathing is a risky behaviour, however, because the fish must approach and break the water surface, which significantly increases susceptibility to predation (Kramer *et al*., 1983). It has been demonstrated that individual MR can drive risk taking to obtain oxygen in fishes (Killen *et al*., 2012; McKenzie *et al*., 2015).

The African sharptooth catfish *Clarias gariepinus* breathes air using a suprabranchial structure known as the arborescent organ. It is a nocturnally active predator, it seeks cover during the day (Willoughby & Tweddle, 1978; Bruton, 1979; Britz & Pienaar, 1992). In captivity, juvenile catfish showed clear diurnal cycles in MR and air-breathing activity, both being much higher at night than during the day (McKenzie *et al*., 2015). In that study, we used respirometry to demonstrate that individual standard MR (SMR, the basal MR of an ectotherm at their acclimation temperature) was a strong driver of rates of oxygen uptake from air. Individuals with high SMR resumed air-breathing more rapidly after a simulated predator attack in their respirometer, evidence that they were bolder. Thus, this appeared to support predictions that individuals with higher MR should routinely take more risks to obtain resources, and be intrinsically more bold (Careau & Garland, 2015; Careau *et al*., 2015; Rupia *et al*., 2016; Krams *et al,* 2017). The time to resume air-breathing after the simulated attack (T-res) appeared to have a binomial distribution, the catfish could be separated into two response groups, ‘bold’ phenotypes with short T-res versus ‘shy’ phenotypes with long T-res. This seemed to indicate that the animals were either bold or shy in this context (Sih *et al*., 2004). The ‘bold’ T-res phenotypes breathed proportionally more air than ‘shy’ phenotypes during the day, when the behaviour was particularly risky due to visibility (McKenzie *et al*., 2015).

In the current study we therefore investigated the general hypothesis that individual SMR and tendency to breathe air would be correlated with major behavioural tendencies such as boldness, activity level and exploration in juvenile *C. gariepinus. We* investigated whether animals with high SMR showed evidence of greater boldness across two contexts, as T-res in a respirometer and as various measures of their behaviour in a novel environment, an open field test (OFT, Archard and Braithwaite, 2011). We expected that correlations between SMR and measures of activity and exploration in the OFT would provide evidence in support of either an ‘increased intake’ or a ‘compensation’ model of how energy metabolism links to these behaviours. We found, once again, evidence that T-res might have a binomial distribution, comprising ‘bold’ individuals with relatively short T-res, and ‘shy’ individuals with relatively longer T-res (McKenzie *et al*., 2015). We therefore investigated whether sorting the catfish into these phenotypes supported our expectations of how SMR would be linked to behavioural tendencies.

## 2. Materials and methods

### 2.1 Animals

Juvenile *C. gariepinus* of unknown sex and a mass of approximately 150 g were obtained from Piscicultura Polettini (Mogi Mirim, SP, Brazil) and transported, by road, to the Department of Physiological Sciences, Federal University of São Carlos (São Carlos, SP, Brazil). There they were maintained in 1 m^3^ tanks supplied with biofiltered well water at 25 ± 1 °C, under a natural photoperiod and fed commercial feed daily at 2% body mass d^-1^, for eight weeks. Animals were then tagged (Passive Integrated Transponder) into the dorsal epaxial muscle under mild anaesthesia (0.1 g l^1^ benzocaine), for individual identification, after which they recovered in routine holding conditions for at least one week before experiments. All experiments were performed at 25 °C.

### 2.2 Metabolic rate and tendency to breathe air

Intermittent stopped flow respirometry (Steffensen, 1989) modified for bimodal fishes (Lefevre *et al*., 2016) was used to measure metabolic rate as oxygen uptake; to evaluate how this was partitioned between water and air, and to derive standard metabolic rate (SMR) as described in (McKenzie *et al*., 2015). Briefly, fish were fasted for 24 h prior to measurements and transferred gently to respirometers without air-exposure, to minimise effects of handling. There were four respirometer chambers, shielded so that spontaneous air breathing behaviours were not influenced by fear of human presence. Fishes were placed in the respirometers in the evening between 18:00 and 20:00 and left undisturbed; respirometry data was collected over 24 h from the following morning.

The setup and methods of bimodal respirometry were exactly as described previously (McKenzie *et al*., 2015; Lefevre *et al*., 2016), with a measure of O_2_ uptake from both phases (aquatic and aerial) collected once every 15 min. Absolute rates of O_2_ uptake from air (*ṀO*_2_a) and water were calculated (in mmol O_2_ kg^-1^ h^-1^) for each 15 min respirometry cycle. They were summed to compute total O_2_ uptake, to then calculate the percentage of this that was obtained from air (%ab). The values for *ṀO*_2_a and %ab were averaged for each individual from 07:00 to 18:00 for daylight (*ṀO*_2_a -day) and from 19:00 to 06:00 for hours of darkness (*ṀO*_2_a -night). Individual standard metabolic rate (SMR) was estimated by the quantile method (Dupont-Prinet et al., 2010; Chabot *et al*., 2016) based upon the 24h of undisturbed measures of total O_2_ uptake (n = 96 per fish), with q fixed at 0.12 such that 12% of values were estimated to fall below true SMR (McKenzie *et al,* 2015; Chabot *et al*., 2016).

### 2.3 Time to resume air-breathing after a fearful stimulus

Following the respirometry, in daylight between 14:00 and 15:00, the screen shielding the setup was lifted for one minute and each fish disturbed by knocking sharply on the lid of their chamber ten times, causing them to retreat to the bottom of the water phase. After this fearful stimulus, the time required for the individual to resume oxygen uptake from air (T-res) was measured in minutes. As previously observed for T-res in this species, data were examined for evidence of a bimodal distribution, comprising a group of ‘bold’ individuals with relatively short T-res, and a second group of ‘shy’ individuals, with relatively longer T-res (McKenzie *et al*., 2015).

### 2.4 Behavioural tendencies in a novel environment

Each individual was studied in an open field test (OFT) as described in Archard and Braithwaite (2011), using a circular opaque dark blue plastic arena (diameter 100 cm • height 80 cm), with water depth 40 cm. The base was marked with a line 12 cm from the sides to delimit edge and centre zones, and the centre zone was divided into 4 equal quarters with two perpendicular lines. The arena was shielded from view by opaque black curtains and illuminated with four 60 W full spectrum daylight bulbs, positioned so as not to be visible to the fish and to leave no shadows within the arena. At the start of a trial an individual was placed, without air-exposure, in a perforated plastic cylinder (diameter 20 cm) in the centre of the arena and left to settle for 5 min. The cylinder was then removed remotely via a pulley, and behaviour was recorded for 10 min, via a webcam positioned above the arena.

Videos were analysed using Etholog v2.2.5 (Ottoni, 2000) to record fish location (edge vs. centre) and movement over the lines on the bottom of the arena. Based upon Archard and Braithwaite (2011), individual boldness was indicated by the time required to choose to return to the centre of the arena (T-centre, in s); the percentage of time spent in the centre, and mean duration (s) of visits to the centre (adjusted for the latency to take cover at the edge when the test began). Individual activity level was indicated by the percentage of time spent moving, and the rate of movement (number of lines crossed / time spent moving (min)). Individual exploratory tendency was indicated by the number of lines crossed / total time (min).

### 2.4 Data analysis and statistics

Statistics were performed with SPSS Statistics vl7.0 (www.ibm.com/software/analytics/spss). SMR showed a significant negative dependence on body mass, by least-squares linear regression of log-transformed data. The residuals of this relationship were therefore used in exploring correlations between individual SMR and the other variables (Killen *et al*., 2011, 2012). None of the other variables showed a significant dependence on body mass and therefore no corrections were applied. Associations among the variables measured in the respirometer and those measured in the OFT were visualised in a principle components analysis (PCA), with the signficance of correlations reported in a matrix of Pearson correlation coefficients. The threshold probability for significance of the multiple correlations was corrected using the false discovery rate (FDR), as described in Pike (2011). Following classification of individuals by their T-res as either bold or shy, all variables were compared between phenotypes by t-test. The level of significance for these tests was α = 0.05.

## 3. Results

The T-res data appeared to be sorted into two groups (McKenzie et al., 2015), either ‘bold’ with a T-res below 95 s or ‘shy’ with a T-res above 115s (Fig 1). The mean values of all variables are, therefore, compared for these two phenotypes in Table 1.

**Figure 1.**
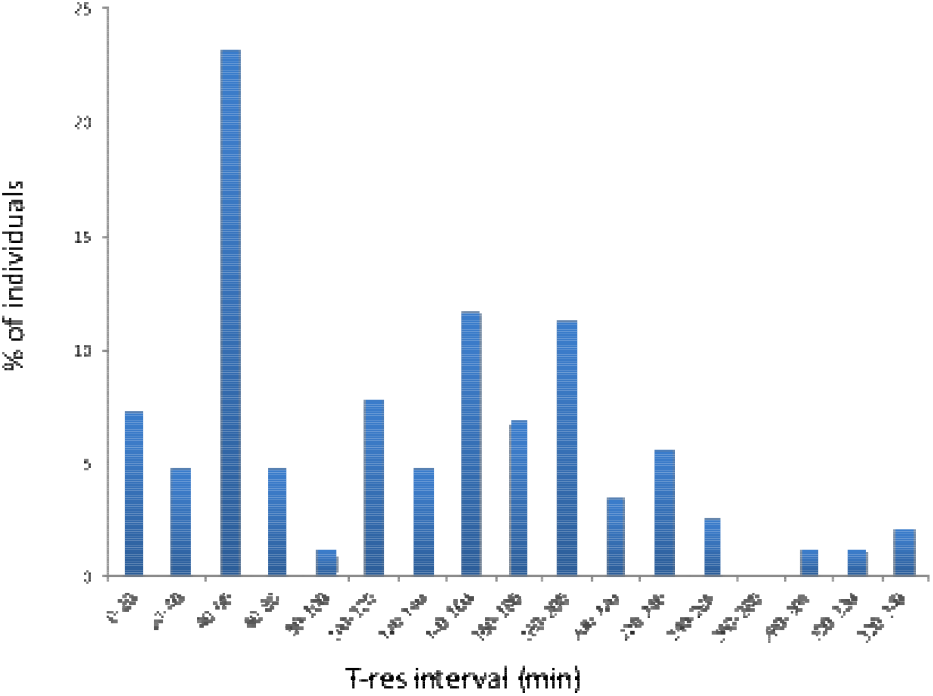
Frequency distribution of T-res phenotypes, showing the bimodal dichocomy into T-res below 85 min or T-res above 115 min.

**Table 1.** Mean (± SE) values for elements of respiratory metabolism and behaviour in an open field test in juvenile *Clariasgariepinus,* overall and when classified by their time to resume air-breathing after a fearful stimulus (T-res) as either ‘bold’ (resumed in less than 80 min) or ‘shy’ (resumed in more than 115 min).

|  | Overall<br>(n = 58) | Bold<br>(n = 27) | Shy<br>(n = 31) |
| --- | --- | --- | --- |
| Mass (g) | 189±75 | 183±72 | 195±78 |
| Respirometric variables |  |  |  |
| SMR | 1.09±0.54 | 1.42±0.11 | 0.80±0.06* |
| T-res | 121±81 | 46.1±3.9 | 186.0±9.5* |
| $\dot{M}O_{2a}$-day | 0.43±0.34 | 0.58±0.08 | 0.29±0.04* |
| $\dot{M}O_{2a}$-night | 0.80±0.62 | 0.93±0.13 | 0.70±0.10 |
| %ab-day | 22±13 | 25.5±2.6 | 19.2±2.1* |
| %ab-night | 35±19 | 35.4±3.6 | 34.2±3.6 |
| OFT variables |  |  |  |
| T-centre | 24.0±7.2 | 21.1±1.1 | 26.5±1.3* |
| Visit duration | 4.1±0.9 | 4.47±0.20 | 3.79±0.11* |
| % T moving | 85±16 | 87.9±2.4 | 82.5±3.3 |
| Move rate | 10.9±3.5 | 9.6±0.4 | 12.0±0.5* |
| Exp rate | 6.9±2.3 | 6.0±0.5 | 7.8±0.5* |

An asterisk indicates a significant difference between bold and shy phenotypes ( $P < 0.05$ T-test). SMR, standard metabolic rate ( $\text{mmol O}_2 \text{ kg}^{-1} \text{ h}^{-1}$ ); T-res, time to resume air-breathing after fearful stimulus (min); $\dot{M}O_{2a}$ , rate of oxygen uptake from air ( $\text{mmol kg}^{-1} \text{ h}^{-1}$ ); %ab, % total oxygen uptake obtained from air; OFT, open field test; T-centre, time to return to centre of arena following commencement of OFT (s); Visit duration, duration of visits to the centre (s); % T moving, % of OFT duration spent moving; Move rate, rate of movement (lines crossed/min moving); Exp rate, exploration rate (lines crossed/min OFT duration).

The first two axes of the PCA only accounted for 49% of the total variation in the data but, nonetheless, revealed associations among variables measured in the respirometer and in the OFT (Fig 2). For example, T-res and an indicator of boldness from the OFT (T-centre) lay close together near axis 1, and were both opposed to SMR along that axis. There were a broad range of significant correlations among the variables (Table 2), with α = 0.020 based upon the FDR (Pike, 2011). Among the respirometric variables, SMR and T-res were, as expected, negatively correlated. Individual SMR was positively correlated with *ṀO*_2_a -day but was not correlated with %ab-day or either measure of aerial respiration at night. T-res, by contrast, was positively correlated with both *Ṁo*_2_a and %ab-day but, once again, there were no correlations with these measures of aerial respiration at night (Table 2). SMR was negatively correlated with a measure of boldness in the OFT (T-centre), positively correlated with % time moving but negatively with the rate of movement (Table 2). T-res was strongly positively correlated with T-centre (Fig 3) and also with rate of movement, in opposition to the effects of SMR (Table 2). There were other correlations of note, for example *ṀO*_2_a -day and *ṀO*_2_a -night were both negatively correlated with % time moving in the OFT, and *ṀO*_2_a -day was also negatively correlated with exploration rate (Table 2).

**Figure 2.**
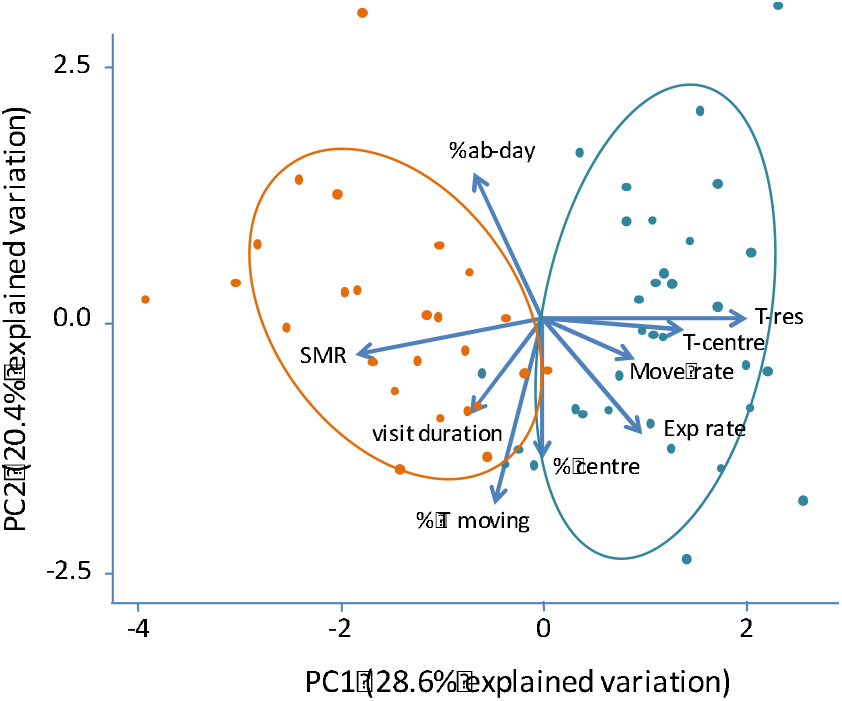
Principle components analysis showing projections of variables on the first two axes, and the projections of the individuals comprising the two T-res phenotypes (bold, orange symbols, vs shy, blue symbols). The ovals denote the 95% confidence ellipses. SMR, standard metabolic rate; T-res, time to resume air-breathing after fearful stimulus; %ab-day, % total oxygen uptake obtained from air in daylight; T-centre, time to return to centre of arena following commencement of OFT; % centre, percentage of total time spent in centre of arena; Visit duration, duration of visits to the centre; % T moving, % of test duration spent moving; Move rate, movement rate; Exp rate, exploration rate.

**Table 2.**
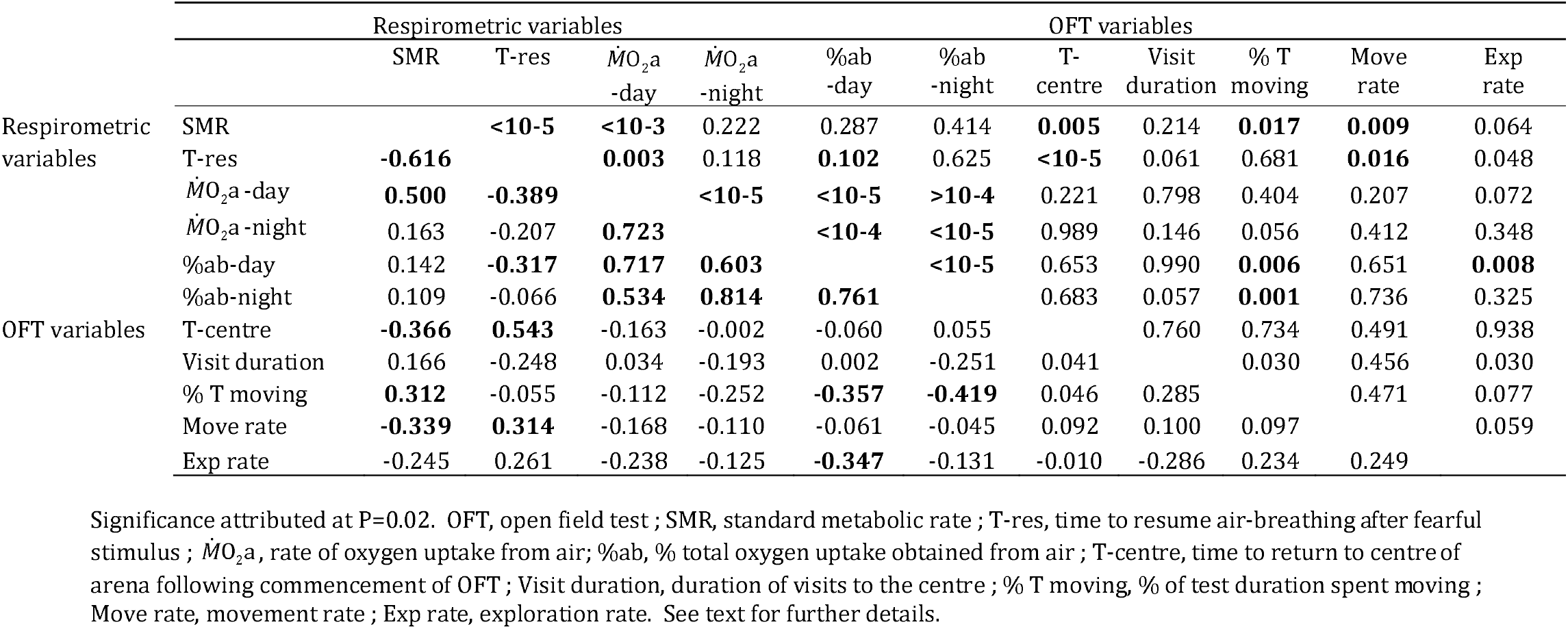
Correlation matrix for respirometric and open field variables. Cells below midline carry Pearson correlation coefficient (R) and those above the midline the associated significance value (P).

**Figure 3.**
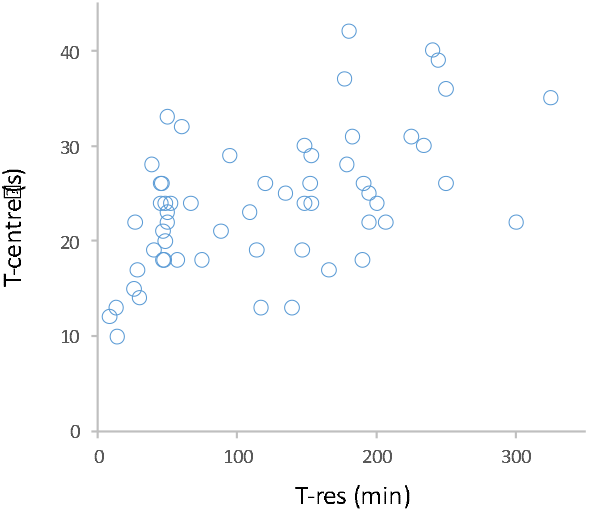
The relationship between time to resume airbreathing after a fearful stimulus in a respirometer (T-res) and time to return to the centre of the arena after commencement of an open field test (T-centre). Dark symbols, ‘bold’ T-res phenotypes; Open symbols, ‘shy’ T-res phenotypes. Although the two variables are highly correlated (P < 0,000001 Pearson correlation for all n = 58 individuals), there is no evidence that T-centre is distributed into two discrete groups in a manner coherent with their T-res phenotype.

When the two T-res groups were projected onto the PCA, they show a clear separation along Axis I (Fig 2). The two groups had significant differences in a number of their respirometric and OFT variables (Table 1). In the respirometer, the bold T-res group had a higher SMR, *ṀO*_2_a -day and %ab-day than the shy, with no differences between bold and shy phenotypes in aerial respiration at night (Table 1). The bold T-res group showed clear indications of boldness in the OFT, with significantly lower T-centre and higher duration of visits to the centre than the shy group (Table 1). By contrast, the shy group had higher rate of movement and rate of exploration (Table 1).

Although T-res and T-centre were very strongly correlated (Table 2, Fig 3), the latter indicator of boldness in the OFT showed no evidence of separation into two phenotypes (Fig 3). The same was true for duration of visits to the centre (data not shown), which differed significantly between the phenotypes (Table 1) but was not correlated with T-res overall (Table 2). Indeed, as displayed in Fig 3, T-res itself seems to be quite continuously distributed.

## 4. Discussion

The data demonstrated that individual metabolic rate and tendency to take risks to breathe air were correlated with behavioural tendencies in an open field context, including boldness, activity and exploratory tendency, in the juvenile catfish. The data provide further evidence of a dichotomy in T-res phenotypes in this species, bold versus shy. Although such a dichotomy was not found in any other behavioural measure, it is nonetheless interesting that, when individuals were sorted into the two putative T-res phenotypes, they differed significantly for their SMR and other variables measured in respirometers and the OFT.

It was noteworthy that individual SMR and T-res were only correlated with air-breathing during the daytime in *C. gariepinus,* which is a nocturnally active species (Bruton, 1979; McKenzie *et al,* 2015). This indicates that these correlations with air-breathing are context-dependent and only visible when surfacing is inherently risky, during the daytime when visible to predators. In the night-time, this perceived risk is presumably relaxed, such that other factors can influence individual air-breathing. The fact that SMR is a driver of individual rates of oxygen uptake from air (McKenzie *et al,* 2015) has a known physiological basis. Air-breathing is a chemoreflex stimulated by oxygen-sensitive receptors in the gills and orobranchial cavity, that sense oxygen in the water and the blood (Smatresk *et al*., 1986; McKenzie *et al*., 1991; Milsom, 2012; Florindo *et al*., 2018). There may be receptors in areas of the venous vasculature where blood oxygen levels reflect rates of oxygen removal by respiring tissues, and stimulate surfacing reflexes more frequently in animals with high tissue oxygen demand (Lefevre et al., 2014; McKenzie et al., 2015; Shelton et al., 1984).

The data also confirmed a negative relationship between SMR and T-res (McKenzie *et al*., 2015) that presumably also has an element of physiological drive, whereby animals with greater oxygen demand were driven to resume air-breathing more rapidly. Nonetheless, the fact that SMR was also negatively correlated to T-centre in the OFT, and that T-res and T-centre were so highly positively correlated, strongly suggest that T-res is also a behavioural manifestation of boldness. A positive correlation between SMR and boldness would be predicted by paradigms to explain why animals show consistent variation in behavioural tendencies and ‘personality’, such as the state:behaviour feedback model (Sih *et al*., 2015) and the pace of life syndrome (Réale *et al*., 2010). That is, these paradigms predict that individuals with higher metabolic rate would exhibit greater boldness in acquiring resources. It has been suggested that this reflects a positive state:behaviour feedback, whereby boldness will be selected for in animals that have a physiological drive to take risks, as this will improve the success of a risky life history strategy. Thus, the high SMR would be a driver of the evolution of boldness (Sih *et al*., 2015). The fact that SMR was not correlated with the proportional extent to which individuals relied on aerial respiration to meet their oxygen demands (the %ab) whereas T-res was, suggests that boldness itself can also be a driver of the tendency to take risks to breathe air (McKenzie *et al*., 2015).

The correlations between SMR and measures of activity in the OFT are notable but difficult to interpret, and can really only be speculated upon. In particular, because questions have been raised about how to interpret behaviour within an OFT (Perals *et al*., 2017). A positive correlation between SMR and the proportion of time that fish were active in the OFT would support the ‘increased intake’ model, whereby animals with high SMR would have a greater metabolic capacity and be able to (or driven to) engage in greater activity (Careau & Garland, 2012). This seems to contrast, however, with the negative correlation between SMR and rate of movement That is, animals with high SMR moved a greater proportion of the time but were doing so at lower speeds, such that actual effort spent on activity is hard to assess. Thus, these data do not clearly support either the increased intake or a compensation model of how energy metabolism links to activity level (Careau & Garland, 2012). Nonetheless, the correlations demonstrate that individual SMR is indeed linked to behavioural tendencies other than boldness, in this air breathing fish.

The mechanisms underlying the negative correlations between %ab, both day and night, and % time moving in the OFT, can also only be speculated upon. That is, it might be expected that animals that breathed more air would be more active rather than less, since activity is a driver of air-breathing in fishes with bimodal respiration (Farmer & Jackson, 1998; McKenzie *et al*., 2012; Lefevre *et al*., 2013, 2014). The negative correlation of %ab-day and exploration rate is counter-intuitive, as it might be expected that animals that tended to take more risks during the day would be more exploratory. These unexpected correlations require further investigation; it should be noted that they are not type II errors linked to the multiple tests, because we corrected our level of significance using the method of False Discovery Rate (Pike, 2011).

The current study provides further evidence of two T-res phenotypes in *C. gariepinus,* with a dichotomy of air-breathing boldness (McKenzie *et al,* 2015). The fact that the two groups were clearly dissociated on the PCA, and showed significant differences in various respirometric and OFT variables, seem to support the existence of a functional dichotomy. The results are consistent with literature predictions that boldness should be associated with high SMR and a risky lifestyle (Réale *et al*., 2010; Careau & Garland, 2012; Sih *et al*., 2015). That is, the bold T-res phenotypes were also bolder in the OFT, had a higher SMR, and breathed more air during the day, both in absolute terms and as a proportion of their metabolic rate. That fact that the bold T-res phenotypes, with their higher SMR, were less active and exploratory in the OFT is consistent with the compensation model (Careau & Garland, 2012). However, the overall correlations did not provide such support for the compensation model, so a clear conclusion cannot be reached with the current dataset It should also be noted that our previous study did not find that the T-res phenotypes differed in their SMR (McKenzie *et al*., 2015), in the current study this was presumably revealed by the larger sample size.

It is puzzling that, although the T-res phenotypes could be assigned to either bold or shy phenotypes in a manner consistent with a previous study (McKenzie *et al,* 2015), none of the measures of boldness in the OFT showed any such evidence. This inevitably raises questions about the validity of the dichotomy in T-res, also because the data may appear dichotomous in a frequency distribution (Fig 1), but seem quite continuous when plotted against another behavioural variable (Fig 3). Nonetheless, finding evidence of a dichotomy in two separate studies is intriguing. This may be linked to how data are interpreted for boldness in an OFT (Perals *et al*., 2017). It is conceivable, also, that the dichotomy is context-dependent and reflects the method of measuring boldness (Sih *et al*., 2004; Thomson *et al*., 2012; Frost *et al*., 2013). Breathing air is a truly risky behaviour that may elicit a predator:prey encounter at the surface. Such risk may promote a dichotomy in bold/shy, as being partially bold or partially shy in such an encounter could be disastrous (Sih *et al*., 2004). On the other hand, individuals in a novel environment may not feel such an immediate risk of predation and may therefore show a spectrum of responses. The literature has no clear consensus on whether bold versus shy tendencies exist along a continuum or are a dichotomy, and how this might depend upon the method of measurement and the context (Sih *et al,* 2004; Thomson *et al*., 2011, 2012; Frost*et al*., 2013).

In conclusion, the data support propositions that animals with elevated metabolic rate, which take risks to acquire resources, are also intrinsically bolder. The relationships of SMR to behavioural tendencies such as activity and exploration were harder to interpret in the current dataset This is consistent with the fact that there is no consensus in the literature about the mechanism that might underlie such relationships. Further research is required to understand whether an apparent dichotomy of boldness might be context dependent, visible by some measures but not others. Air-breathing fishes are useful models to explore relationships between physiology and behaviour (McKenzie *et al*., 2015; Killen *et al*., 2018; Pineda *et al*., 2020), future research should confirm that behavioural tendencies are temporally stable and therefore constitute elements of personality (Conrad *et al,* 2011; Niemelä & Dingemanse, 2018). These species could then be used to test hypotheses about how energy metabolism links to personality.

## 5. Acknowledgements

This study was funded in part by the Brazilian National Council for Scientific and Technological Development – CNPq (313621/2013-6) and the State of São Paulo Research Foundation – FAPESP (2015/22326-0) to DJM; the Fundação Coordenação de Aperfeiçoamento de Pessoal de Nível Superior – CAPES (Master fellowship to FRB), and a Natural Environment Research Council Advanced Fellowship (NE/J019100/1) to SSK. The authors are grateful to Cesar Polettini, of Piscicultura Polettini, for donating the catfish.

